# Using the novel priority index in prioritizing the selection of inland water bodies for site-based fish species conservation

**DOI:** 10.1101/2020.11.12.380618

**Authors:** Anthony Basooma, Herbert Nakiyende, Mark Olokotum, Winnie Nkalubo, Laban Musinguzi, Vianny Natugonza

## Abstract

Freshwater ecosystems occupy <1% of the Earth’s total surface area but provide an array of ecosystem services. However, these ecosystems are threatened by multiple stressors, including overexploitation, infrastructure developments, habitat alteration, and alien species introductions. The magnitude of these threats varies in different water bodies, requiring site-based conservation actions. In this paper, we aimed at developing a priority index (CPIw) that can be used to inform conservation managers in prioritizing the selection of a waterbody for site-based fish conservation purposes. We used data on distribution, diversity, and conservation status of fishes of Uganda, which were retrieved from the Global Biodiversity Information Facility (GBIF) and International Union Conservation for Nature (IUCN) databases. In the index, we incorporated the species richness, surface area of a waterbody, species rarity, and species IUCN status. A total of 288 fish species were recorded in 81 waterbodies (7 large lakes, 37 small lakes, and 37 rivers). Of these species, 110 were only found in large lakes, followed by rivers (19) and small lakes (6). Despite the higher species richness in large lakes relative to small lakes, the latter recorded significantly higher CPIw compared with the former (t = −2.8, df = 30, p-value = 0.008, d=0.7). This observation is consistent with the expectation, given the low ecological substitutability for the species and higher levels of exposure to human-induced threats in small water bodies compared with large systems. Therefore, we suggest that in situations where resources are limiting, small water bodies need to be given much attention, although we do not suggest ignoring water bodies with low CPIw values.

## Introduction

Freshwaters constitute ~3% of the Earth’s water (WWF 2020) and occupy <1% of the Earth’s surface area (Garcia-Moreno et al. 2014), but is habitat to about 40% of all described fish species (Lundberg et al. 2000). The benefits of these freshwater resources to society are immense. For example, 21 largest lakes in the world provide about 1.3 million tons of fish annually, 62.5 GW of hydropower, 5 billion m^3^ of potable drinking water, and 815 million m^3^ of water for irrigation (Sterner et al. 2020). Also, freshwater systems replenish estuarine, oceans, and seas with nutrients and water (Matthews 2016); for example, Nile River deposits 1318 and 212 kg km^−2^yr^−1^ of nitrogen and phosphorus, respectively, into the Mediterranean Sea (Yasin et al. 2010).

Freshwater species are more susceptible to human-induced threats such as climate change, pollution, habitat alteration, overexploitation, and alien species introductions compared with marine and terrestrial species (Darwall et al. 2018; WWF 2020). For example, while marine and terrestrial species have declined by 39% in the last 50 years, freshwater counterparts have reduced by 76% higher than the global average of 52% (WWF 2014). Freshwater fish species might be the most threatened vertebrates assessed by IUCN (Reid et al. 2013). The species are affected by the high levels of exploitation to support about 158 million people worldwide who derive animal protein from freshwater fish species (McIntyre et al. 2016).

Approximately 500 fish species are reported in all water bodies in Uganda (NEMA 2007; Natugonza & Musinguzi, 2020). However, several non-native fishes, including Nile perch (*Lates niloticus),* Nile tilapia (*Oreochromis niloticus*), redbelly tilapia (*Coptodon zillii)*, blue-spotted tilapia (*Oreochromis leucostictus*), and redbreast tilapia (*Coptodon rendalii*) were introduced into various lakes and rivers within the Victoria and Kyoga lake basins (Kishe-Machumu et al. 2018). These introductions especially for Nile perch coincided with the collapse of most native fish species (Ogutu-Ohwayo 1990). In Lake Victoria, ~300+ haplochromine cichlids were extirpated (Kaufman 1992; Ogutu-Ochwayo 1990), and similar destructive ecological changes were observed in lakes Kyoga and Nabugabo (Ogutu-Ohwayo 1990; Chapman et al. 1996). This loss in haplochromine cichlids is believed to be the worst vertebrate species extinction observed in recent times (Kaufman 1992), placing Nile perch among the 100 top worst alien invasive species in the world (Lowe et al. 2000). Other native species such as Singidia tilapia (*Oreochromis esculentus*) and Ningu (*Labeo victorianus*), which previously dominated in fish catches from Lake Victoria and its affluent rivers (Kudhongania et al. 1992) are currently classified as critically endangered (IUCN 2020). The catfishes, including Semutundu (*Bagrus docmak* Forsskål, 1775), Lake Victoria deepwater catfish (*Xenoclaris eupogon*), *Clarias* spp.,*Synodontis* spp., and Silver catfish (*Schilbe intermedius*) were also affected by Nile perch establishment in Lake Victoria (Goudswaard & Witte 1997; Balirwa 1998; Balirwa et al. 2003). Riverine species were mostly affected by overexploitation and habitat degradation, while the native tilapines declined mostly through interspecific competition and hybridization (Cadwalladr 1965; Kudhongania et al. 1992).

The reduction in fish species diversity in Uganda has led to numerous studies focussing on the species diversity, abundances, distribution, taxonomy, and biology of the remnant species to facilitate their recovery and reduce further extinctions (Witte & Van Oijen 1990; Kaufman & Ochumba 1993; Ogutu-Ohwayo 1993; Ogutu-Ohwayo et al. 1999; Mbabazi et al. 2004). In particular, small lakes, swamps, rivers, streams, and wetlands were documented as the main structural refugia for these fishes (Ogutu-Ohwayo et al. 1999; Mwanja et al. 2001; Chapman et al. 2002; Mbabazi et al. 2004; Balirwa et al., 2003; Wakwabi et al. 2006; Olwa et al., 2020). However, these studies were waterbody-specific with limited information to rank the waterbodies for site-based conservation given the limited available resources. Analysis of the distribution of fishes at a broader scale has been limited in the past due to the paucity of data, which have been scattered in many research institutions in unusable formats. Recently, substantial amounts of data on the occurrence of fishes for Uganda have been made available through the Global Biodiversity Information Facility portal (GBIF) (GBIF, 2020). This study aims to develop a Conservation Priority Index (CPIw) for inland water bodies to prioritize their selection for site-based fish conservation, especially when resources are limiting. We use data on the distribution, diversity, and conservation status of the fish species in Uganda, which are freely accessible through GBIF and IUCN databases.

## Methods

### Data acquisition and processing

We retrieved occurrence records from two fish classes (Actinopterygii and Sarcopterygi) in Ugandan water bodies from GBIF online data repository (GBIF 2020). We used the *occ download get* function in *rgbif* package to retrieve data (Chamberlain et al. 2020). Except for genera *Astatoreochromis and Pseudocrenilabrus*, we changed all other haplochromine cichlids genera to *Haplochromis*to conform with FishBase nomenclature (Froese & Pauly 2019), which is based on Oijen (1996). Thus, names such as *Astatotilapia nubila*were changed to *Haplochromis nubilus*; *Schubotzia eduardiana* to *H. eduardianus*; and *Astatotilapia* pallida to *H. pallidus*. Occurrences that were outside the geographic range described in FishBase, and whose identity could not be verified based on recent survey data, were discarded. We also excluded all occurrences without complete scientific names (genus and specific epithet), e.g., *Haplochromis* sp. and *Oreochromis* sp. Occurrences with unknown and incorrect water bodies were excluded, e.g., all records of *H. eduardii* that were recorded in Lake Albert in the GBIF datasets were discarded, as the species is endemic to Lake Edward (Froese & Pauly 2019). For occurrence records without a named waterbody of origin but with coordinates were determined based on the GPS coordinates. We used habitat descriptions, verbatim locality, and location remarks to identify the waterbody of origin. We also discarded all occurrences from manmade water bodies, such as ponds, tanks, and aquarium. Lakes Salisbury and Kasudho were changed to Bisina and Kasodo, respectively, to conform to the current names and avoid duplication of records for the same lakes. We categorized lakes <200 km^2^ as small lakes and >200km^2^ as large lakes, resulting in 7 large and 37 small lakes (Appendix S1 and S2). After preliminary processing of the data, a total of 14,452 occurrences records were retained for further analysis.

### Conservation priority index formulation

We retrieved the conservation status of each species from the International Union for Conservation of Nature (IUCN) Redlist database (www.iucnredlist.org/). The species are classified as data deficient (DD), least concern (LC), critically endangered (CR), near threatened (NT), endangered (EN), and not evaluated (NE) (IUCN 2012). We used waterbody surface area, species richness, IUCN statuses, rarity, and scaling constant to develop the conservation priority index (CPIw), based on the formula:

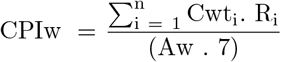

where, for species *i* per unit surface area of a waterbody, Cwt_i_ is the species weight based on its IUCN status (i.e., CR (5), EN (4), VU (3), NT (2), LC (1)); n is the number of species in a particular waterbody; Aw is the total surface area of the lake, and Ri is the frequency of occurrence of a species in the water bodies (i.e., if a species occurred in one waterbody, then a weight of 5 was assigned, 2-3 water bodies (weight 4), 4-5 water bodies (weight 3), 6-10 water bodies (weight 2), and >10 water bodies (weight 1)). The value 7 is a scaling constant, indicating the total number of IUCN categories (IUCN, 2012). Note that fish species that were registered as NE and DD in the IUCN database were assigned a Cwt_i_ of 5 on the basis that such species can go extinct unnoticed and, therefore, should be considered in the same category as CR species (IUCN 2012). The surface area for each lake was obtained from the literature (Burgis & Symoens 1987; Vanden Bossche & Bernacsek 1990; Ogutu-Ohwayo et al. 1999; Schofield & Chapman 1999; Olowo et al. 2004). We used Google Earth to approximate the surface area for lakes Gawa, Kabaleka, Wamala, Nakabale, Owapet, Kirimira, and Kabaka because it was not found in the literature. The surface area of Lakes Natuali, Chankaranga, Okurachere, Kasunju, Nkuruba, and Mutabyo could not be determined from both literature and Google Earth, and thus we could not calculate their CPIw values.

### Data analysis

For each water body category (small lakes, large lakes, and rivers), we determined the species richness and the total number of species in IUCN status categories. We used a non-parametric Kruskal Wallis test to assess the mean differences in the species richness and IUCN categories among waterbody categories. Post hoc multiple comparisons were conducted with a Dunn’s test to determine the statistical differences between the waterbody categories. We generated a species accumulation curve for the waterbody to assess if most of the species found in the data. We determined the rarity of a particular species by summing the frequency of occurrence in the water bodies where it was found. We used the Bray Curtis dissimilarity measure to compute the ranks for water bodies and species and later used a non-metric multidimensional scaling (nMDS) to visualize the species and water bodies in 2-D ordination space. After, we performed an analysis of similarity (ANOSIM) to determine the statistical differences among waterbody categories. A similarity percentage analysis (Simper) was used to evaluate the contribution of the species to the dissimilarities between waterbody categories.

For the index, we log-transformed the CPIw values and used a Shapiro-Wilk and Levene tests to examine for normality and equality of variance, respectively. After, we used a parametric Welch 2-Sample t-test to evaluate the differences between the mean CPIw values for large and small lakes. We processed data with predefined functions in R (including *specaccum, diversity, metaNMDS, simper, anosim,*) of the *Vegan* package (Oksanen et al. 2019), *dunn test* in Dunn package (Dinno, 2017).

## Results

### Species composition and rarity

A total of 288 species were recorded from 81 water bodies (i.e., 7 large lakes, 37, small lakes, and 37 rivers) (Fig 1 & Appendix S2). Of the 288 species, 163 were haplochromine cichlids and 125 non-haplochromine. The species accumulation curve increased at a low rate after 20 water bodies (Fig 2). All haplochromine cichlids were recorded in 45 water bodies compared with 75 for non-haplochromines. Species richness was highest in large lakes dominated by Lake Victoria with 175 species followed by Albert, Kyoga, and George (Fig 1, Appendix S2). Kruskal Wallis test showed that waterbody categories differed significantly in mean species richness, and the IUCN status categories (Table 1). However, the nature of a waterbody had a weak effect on the variation in species richness and IUCN categories (Table 1). Post hoc multiple comparisons showed that large lakes were significantly different from rivers and small lakes for all IUCN categories and species richness. However, except for the vulnerable category, rivers and small lakes were not significantly different (Table 1).

**Table 1.**
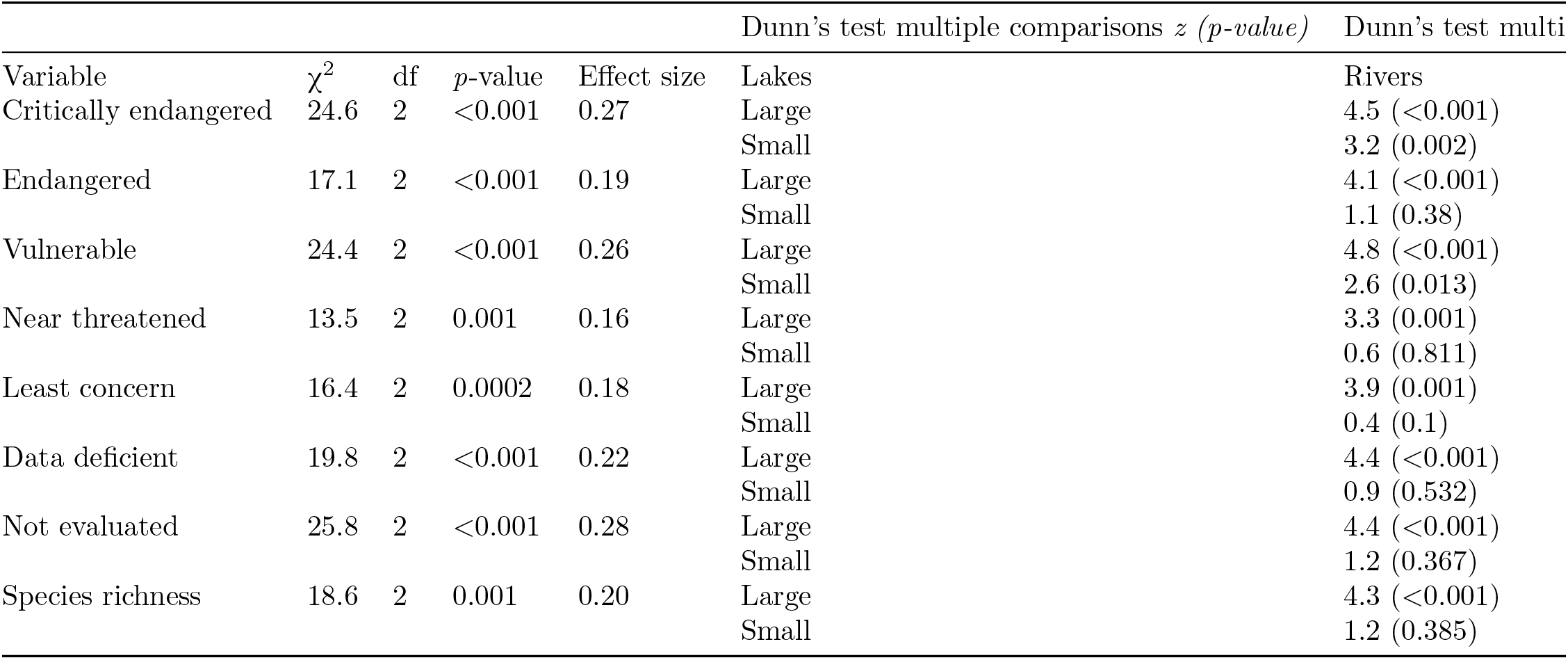
Kruskal Wallis analysis of variances for IUCN categories and species richness followed by Dunn’s multiple comparison test between waterbody categories

**Figure 1.**
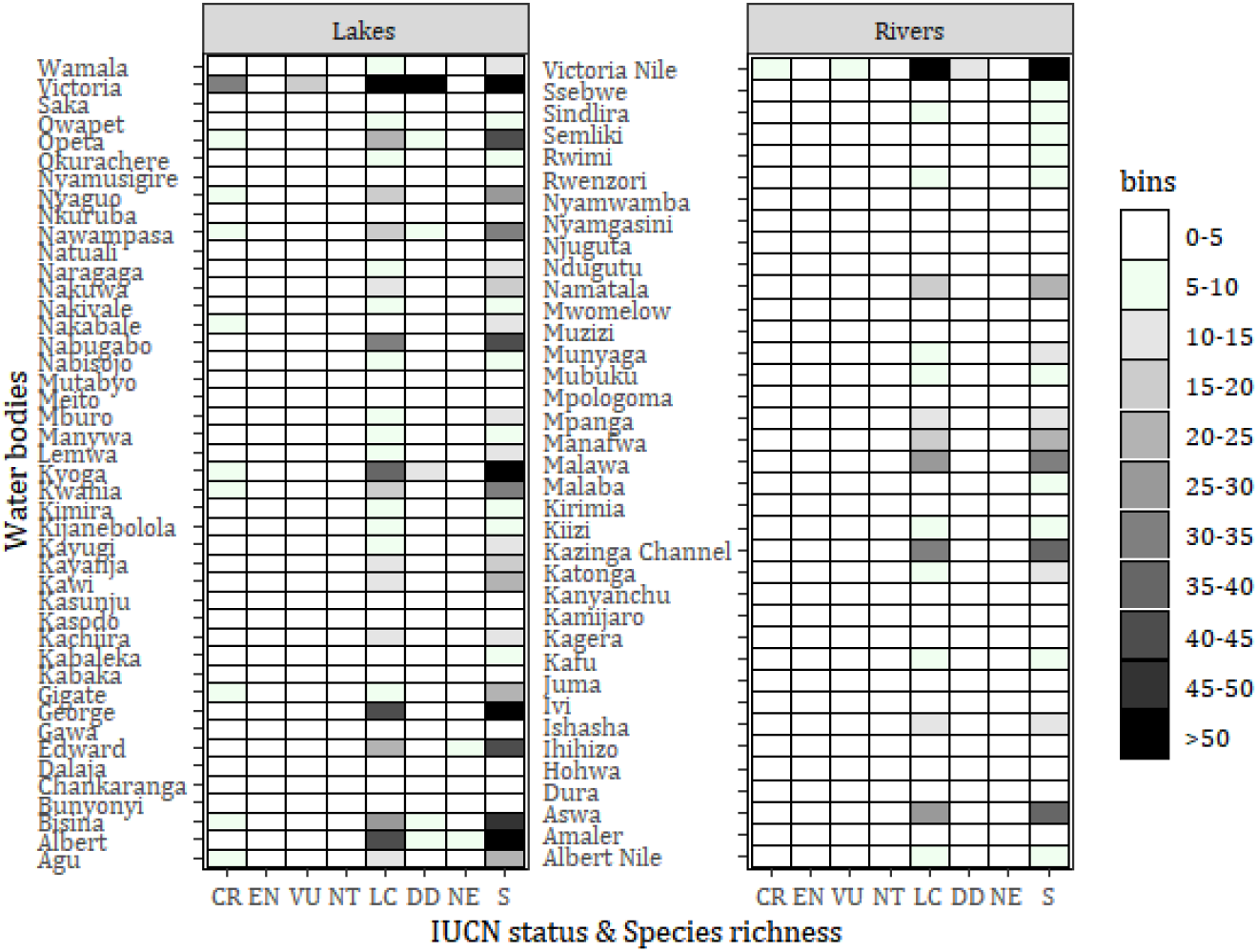
Species composition in water bodies according to IUCN status and richness.

**Figure 2.**
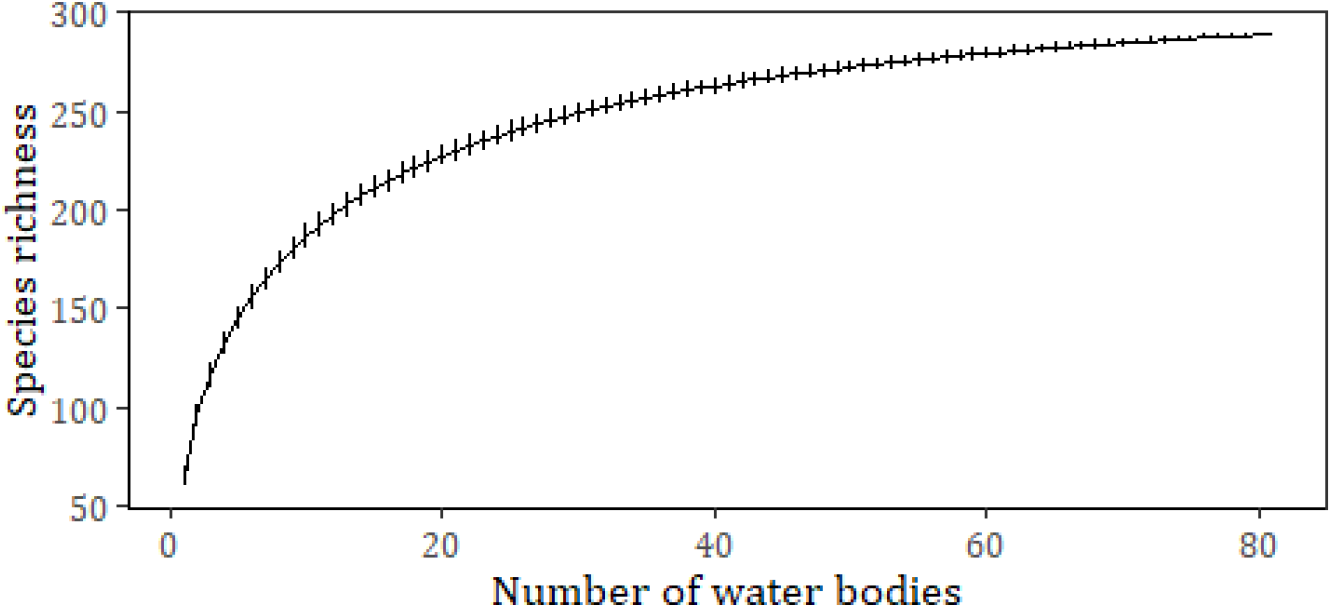
Species accumulation curve for fish species from 81 water bodies in Uganda

Of the 163 haplochromine cichlids, 85 species were found in utmost one waterbody compared with 31 non-haplochromine species (Fig 3). Lake Victoria had 71 rare species, where 11 were critically endangered haplochromine cichlids and one non-haplochromine species (*Xenoclarias eupogon*) (Appendix S3). Two near threatened species (*H. labiatus and H. oregosoma*) were only found in the interconnected system of lakes George, Edward, and Kazinga Channel.*Clarias gariepinus* was found in 42 water bodies followed by *Enteromius kerstenii*, *O. niloticus* (33), and*Protopterus aethiopicus* (32). *Astatoreochromis alluaudi* was recorded in 30 water bodies, *H. nubilus* (28), *H. lividus* (16), *H. argenteus* (14), *H. phytophagus* (13),*H. obesus* (12), and *H. parvidens* in (11).*Oreochromis esculentus* was observed in 19 water bodies, *O. variabilis* (15), and *L. victorianus* (8). Based on IUCN categories, no vulnerable non-haplochromines was found in Lake Victoria; however, 8 of the 25 vulnerable haplochromine cichlids were only recorded in the lake (Appendix S3). *Labeobarbus ruwenzorii* and*L. alluaudi* were only found to Lake Edward and rivers Ivi, Rutushuru, Rwimi, Sebwe, Rwenzori, and Mubuku). *Nothobranchius taeniopygus* and *Synodontis macrops* were only found in Aswa River. *H. melanopterus* was recorded in Lake Kasodo whereas*H. commutabilis* and *H. ampullarostratus* were only found in Lake Kachiira (Appendix S3). Of the endangered fish species,*Astatotilapia desfontainii* was only recorded in lakes Bisina, Victoria, and Victoria Nile; *H. simpsoni* was only found in lakes Kyoga, Nabugabo, and Kayanja, while *Lates macrophthalmus* was only recorded in Lake Albert. *H. beadlei*, a critically endangered species, was found in lakes Nabugabo and Victoria, and *H. granti* in Lake Victoria and River Kagera.

**Fig 3.**
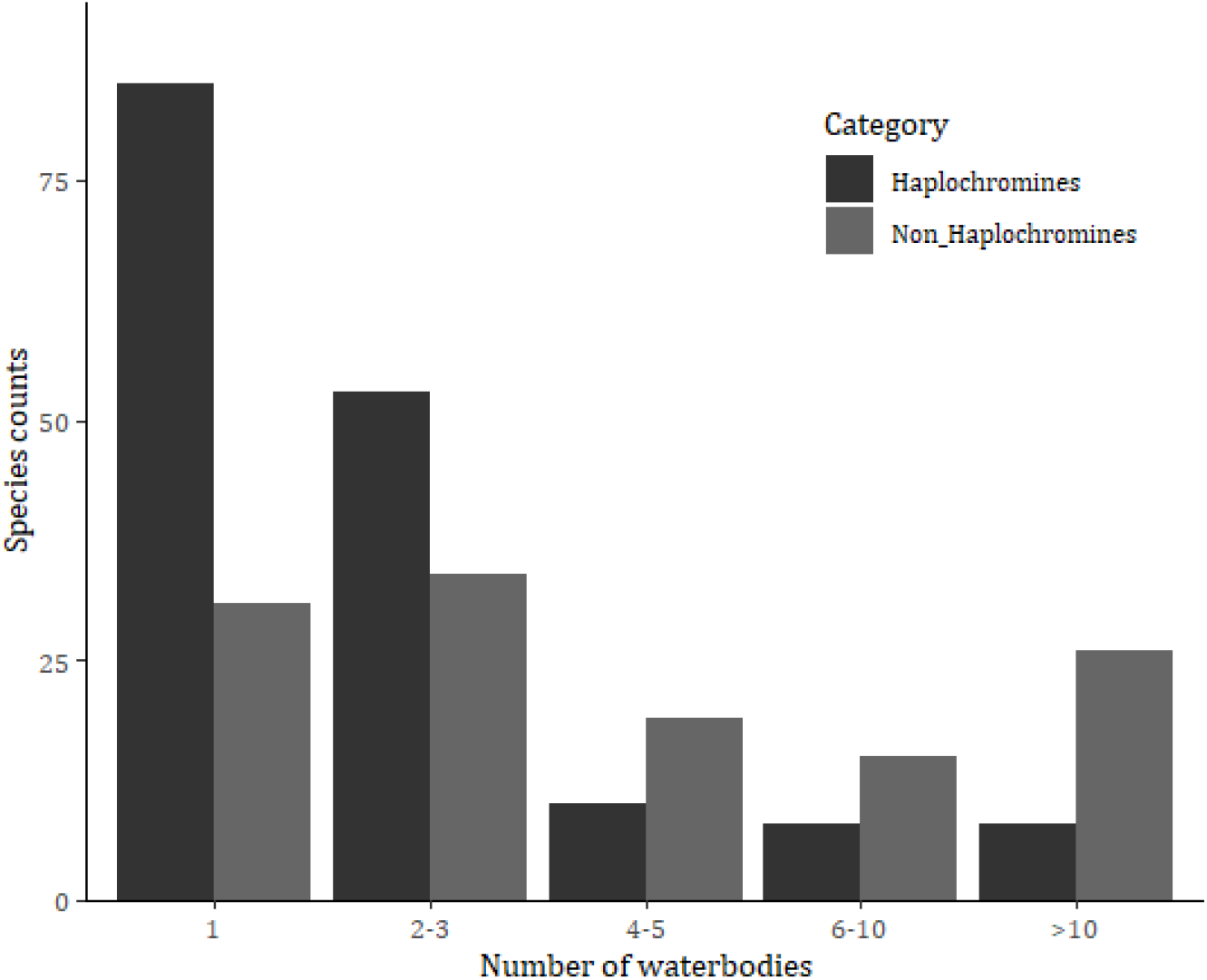
Rarity of species in the different water bodies in Uganda

Among the 3 waterbody categories, large lakes recorded the highest number of non-shared species (110), followed by rivers (19), and the small lakes (6) (Fig 4). The species richness in waterbody categories differed significantly (Kruskal Wallis: χ^2^= 18.6, df =2, p=0.001, η^2^ = 0.2). The nMDS had a stress level of 0.14, with large and small lakes closely clustered except rivers (Appendix S4). Haplochromine cichlids were closely clustered, except*H. latifasciatus, H. victoriae, A. alluaudi, H. nubilus, and H. schubotzi*, while non-haplochromine species were widely distributed (Appendix S5). Analysis of similarity (ANOSIM) among waterbody categories showed significant differences (R=0.24, p<0.001). Similarity percentage (simper) analysis showed that large lakes differed from small lakes (93.5%) and rivers (97.5%), while small lakes differed from rivers by 96.2%. *Lates niloticus* and *O. niloticus* contributed the highest average percentage variation between large lakes and rivers. Similarly, *L. niloticus* had the highest contribution between large and small lakes (Appendix S6). In contrast,*A. alluaudi* and *C. gariepinus* had the highest contribution to the variation between rivers and small lakes.

**Figure 4.**
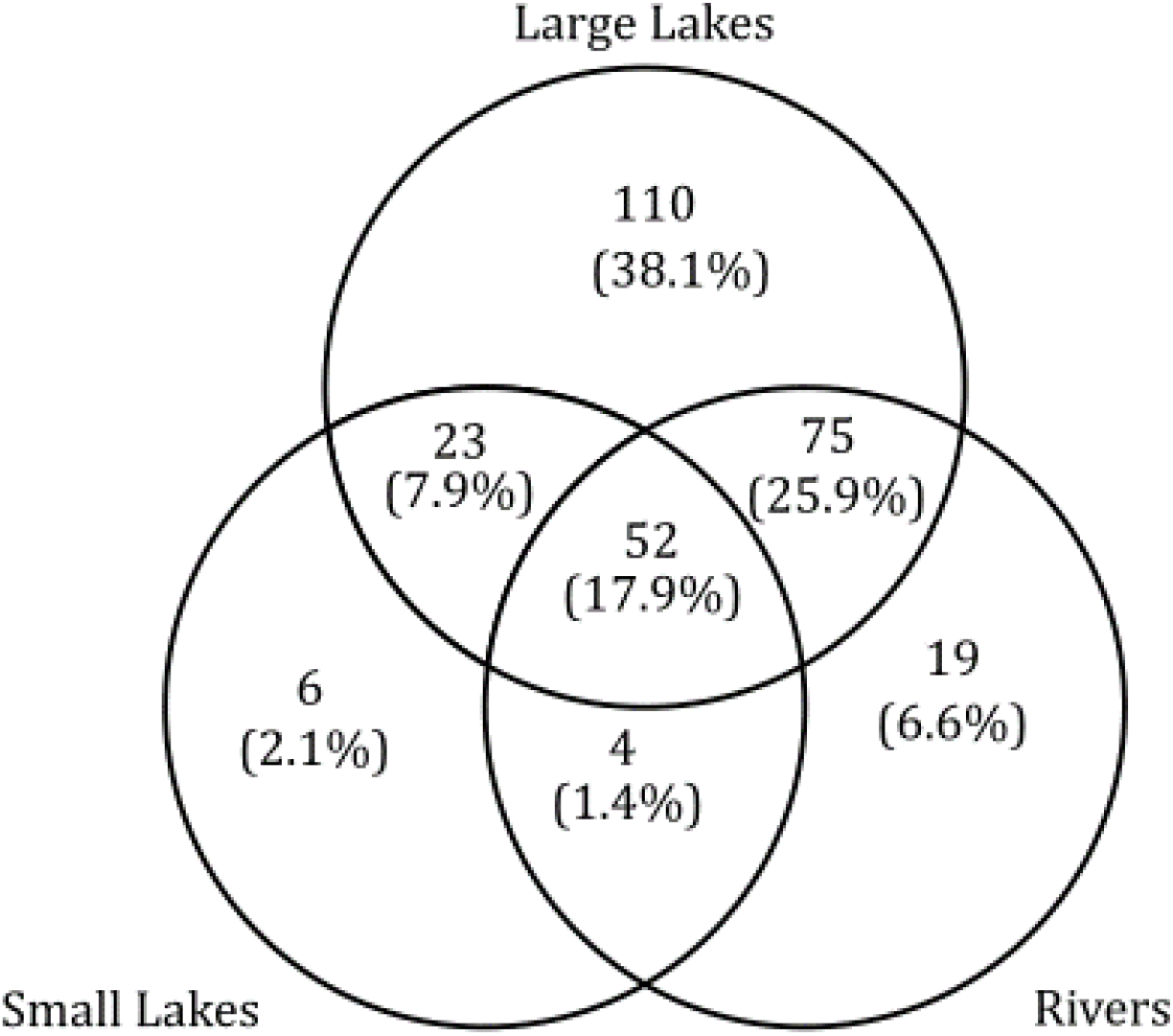
Fish species composition in the three water body categories

### Conservation Priority Index (CPIw)

CPIw was computed for 38 lakes, where 7 were large lakes and 31 small lakes. Small lakes had a mean CPIw of 2.4 (SD 4.6) compared with 0.05 (SD 0.04) for large lakes. A significant difference between the mean CPIw value for large and small lakes was observed (Welch 2-Sample t-test: t = −2.8, df = 30, p-value = 0.008, d=0.7). The surface area of the lakes had a large effect on the CPIw values. Highest CPIw values were recorded for lakes Manywa followed by Kayanja, Gigate, Agu, Naragaga, Kawi, and Nabugabo (Fig 5 & Appendix S7). Low values were mostly recorded for large lakes and highest for small lakes.

**Figure 5.**
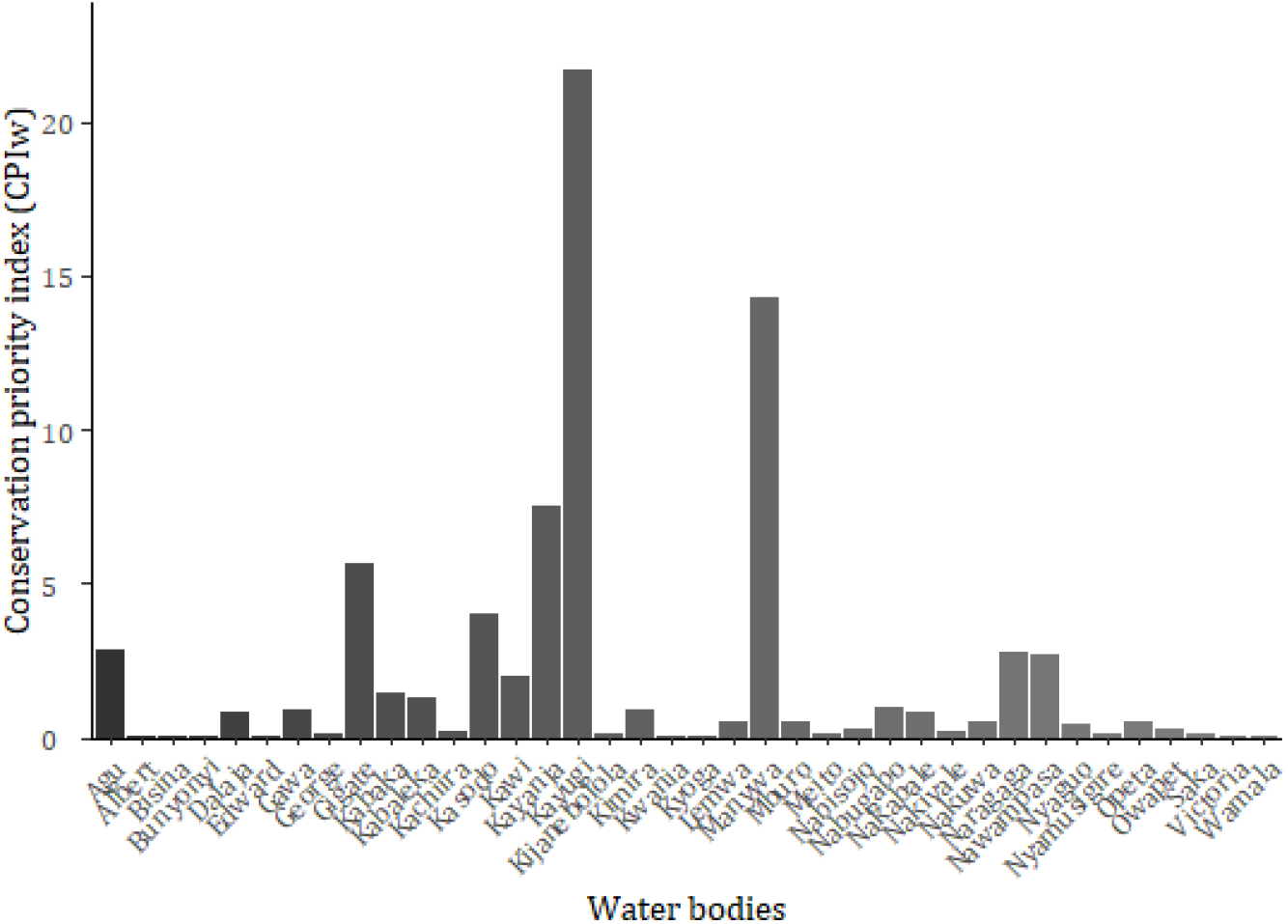
Conservation Priority Index (CPIw) for evaluated water bodies in Uganda.

## Discussion

### Species composition, rareness, and distribution

The total species count recorded was comparable with the FishBase overall estimate of 278 described fish species in Uganda (Froese & Pauly 2019). Also, the species accumulation curve approached the asymptote, suggesting that most of the species were considered in the analysis (Gotelli & Colwell 2001). Large lakes had higher species richness and more rare species compared with rivers and small lakes. Differences in species richness and rarity among water bodies may be attributed to geomorphological, abiotic, and biotic factors (Brown et al. 2007). Also, isolated inland water bodies favor rapid allopatric speciation and adaptive radiation (Basiita et al. 2018), which exposes the species to different evolutionary pressures. For example, along the Victoria Nile, Murchison Falls hinders fish migration to Lake Kyoga from Albert (Basiita et al. 2018). The falls along River Semiliki prevents fish passage to Lake Albert from Edward (Acere & Mwene-Beyanga 1990). Similarly, between Lake Victoria and Kyoga have been separated by the Owen and Bujagali falls (Basiita et al. 2018), and a sandbar between Lake Nabugabo and Victoria (Stager et al. 2005). These biogeographical barriers may have led to allopatric speciation; for instance, Lake Nabugabo which was once connected to Lake Victoria had five endemic species (Ogutu-Ohwayo 1993; van Alphen et al. 2004). At the species level, *L. niloticus* from lakes Kyoga and Victoria were genetically different (Basiita et al., 2018) possibly due to manmade barriers between the two lakes, which has impeded gene flow (Basiita et al. 2018).

High species richness and rarity are expected in habitats of long geological age, geographically isolated, and where species are prone to speciation (Strayer 2013). Also, according to the biogeographical principle, large areas have more species (Rosenzweig 1995). In this study, the high species richness and rareness observed in Lake Victoria is not surprising, given its surface area, explosive speciation and adaptive radiation of haplochromine cichlids because of hybridization, allopatric, microallopatric and sympatric mechanisms (van Alphen et al. 2004; Meier et al. 2017). A similar trend of high species richness with increased lake area was observed (Amarasinghe & Welcomme 2002). In large water bodies, species can shift to favorable habitats, for example, *Squalius lucumonis* and *Telestes muticellus* that migrated upstream in the Mediterranean Rivers due to climate warming (Carosi et al. 2019). The level of rarity of recently described species could not be determined in the available data. For example, the distributions of *H. akika* in Lake George (Lippitsch 2003) and*H. katonga* in River Katonga (Schraml & Tichy 2010) are not well understood except where the specimen was obtained. Also, haplochromine cichlids are mostly lumped as haplochromines (Marshall 2018), which may have accounted for the absence of certain species in particular water bodies.

For rivers, habitat degradations and manmade obstructions mostly damming have affected fish species migration, feeding, and recruitment patterns (FAO 2001). For instance, only 70 of the 177 large rivers in the world are free from damning (WWF, 2020). In Uganda, along the Upper Victoria Nile, three dams were constructed. These obstructions affected species richness and gene flow along the river (Basiita et al., 2018). For example, the stocks of *H. simotes* and *L. victorianus* have been affected by dams along the Upper Victoria Nile (Sayer et al. 2018). Similar effects of damming have been reported along Yangtze and Mekong (Dugan et al. 2010; Yi et al. 2010).

### Conservation Priority Index (CPIw) for inland water bodies

Biological metrics including indices, indicators, or targets have been used by managers and decision-makers to prioritize areas for conservation (Tognelli et al. 2019; Linke et al. 2011). These metrics may include prioritizing areas with threatened species (Kirkpatrick 1983), threatened species affected by climate change (Tognelli et al. 2019), or fixed percentages of an area (Linke et al. 2011). However, some indices such as species richness do not incorporate the component of complementarity of the areas in question, thus, highly-ranking areas with the same species (Kirkpatrick, 1983). Shannon Weaver diversity index does not distinguish habitats with the same species evenness and richness (Omayio & Mzungu, 2019). Most priority indices are computed for terrestrial ecosystems or specific for a particular region, thus, cannot be extrapolated to other systems (Brum et al. 2017). For example, the Forest Conservation Priority index (FCPI) (de Mello et al. (2016) used the area and shape of the forest without considering the species conservation status. The Cave Conservation Priority Index (CCPi) (Souza Silva et al. 2014) did not consider the species or their conservation status but species richness, distribution, and impact weights. In the species conservation importance index, the species conservation status and rarity were considered (Halmy & Salem, 2015), but the index applied to terrestrial plants but not an aquatic ecosystem. Thus, these priority indices cannot be used to rank water bodies for site-based priority conservation.

In most instances, biodiversity measures such as species richness are usually higher for large water bodies, which would mean they are prioritized for conservation. Indeed, this study also showed higher species richness for large lakes compared to small ones. However, the novel conservation priority index (CPIw) was significantly higher for small lakes compared with large lakes. This observation is consistent with expectation, given the low ecological substitutability for the species and higher levels of exposure to human-induced threats in small water bodies compared to large systems. For a system such as Lake Victoria, with a vast habitat heterogeneity, fish species can easily seek refugia in other habitats (Seehausen et al. 1997; Chapman et al. 2003), which may not be possible in a small water body. The index showed that, for example, Lake Gigate with a surface area of 1.7 km^2^, but with 4 critically engendered haplochromine cichlids (*H. latifasciatus, H. obesus, H. parvidens,* and *H. argenteus*) could be prioritized for conservation ahead of a large system with higher species richness. The likelihood of the species getting extinct in Lake Gigate is eminent if a similar magnitude of stress is applied to both lakes. However, the index does not imply that other water bodies with low CPI should not be monitored, but it may allow a conservation manager or decision makers to rank water bodies for urgent intervention, especially if the resources are limiting. Because conservation interventions should address social needs (Linke et al., 2011), large waterbodies, which are usually productive would be difficult to fully conserve. The index should, therefore, be adopted as a rapid metric measure to rank water bodies to enable prioritizing them for conservation. Further, if the size and species in habitats in the waterbody are known, the index can be downscaled to a habitat level. However, index could not be applied on rivers because most of them are dammed or obstructed, creating distinctive habitats along the river.

## Supporting Information

The map large lakes in Uganda (Appendix S1), Waterbody species richness and IUCN status (Appendix S2), Rare species (Appendix S3), Ordination plot for waterbodies (Appendix S4). Ordination plot for species (Appendix S5), Simper analysis plot (Appendix S6), Waterbody Conservation priority index values (Appendix S7)

